# Simulated microgravity accelerates alpha-synuclein aggregation and induces oxidative stress in an *in vitro* Parkinson’s disease model

**DOI:** 10.1101/2024.10.25.620204

**Authors:** Veronica Lentini, Giuseppe Uras, Alessia Manca, Mohammed Amine El Faqir, Sara Lucas-Del-Pozo, Nicola Deiana, Giacomo Cao, Anthony HV Schapira, Antonella Pantaleo

**Affiliations:** Department of Biomedical Science, University of Sassari, Viale San Pietro, Sassari, 07100, Italy; Department of Clinical and Movement Neurosciences, UCL Queen Square Institute of Neurology, London, UK; Department of Mechanical, Chemical and Materials Engineering, University of Cagliari, Piazza d’Armi, 09123 Cagliari, Italy

## Abstract

Parkinson’s disease (PD) is a neurodegenerative disorder characterized by the accumulation of alpha-synuclein aggregates and progressive neuronal loss in the substantia nigra, with aging being its primary risk factor. The current available models to study PD mechanisms are largely relying on genetic mutations to recapitulate PD typical hallmarks, such as increased alpha-synuclein aggregation. However, they do not model the aging features associated with the disease.

Microgravity, a condition experience by astronauts during space missions, is known to induce ageing-like modifications on both systemic and cellular physiology.

To replicate the aging-related stress observed in PD patients, we exposed SH-SY5Y and 3K-SNCA mutant cell lines to simulated microgravity.

Our findings revealed that simulated microgravity enhanced PD alterations, with a significant increase in misfolded and phosphorylated a-syn. This was accompanied by heightened oxidative stress, as evidenced by increased levels of reactive oxygen species, without a sufficient antioxidant response. These results suggest that simulated microgravity effectively mimics and accelerate the stress associated with aging in PD cell models, regardless of the presence of PD mutation. This study highlights the potential of simulated microgravity as a tool for investigating aging processes in neurodegenerative diseases.

## Introduction

Parkinson disease (PD) is the second most common neurodegenerative disorder and is characterized by both motor and non-motor clinical features [1–3]. PD is defined pathologically by the presence of alpha-synuclein (a-syn) aggregates in Lewy Bodies and neurites with associated loss of pigmented dopaminergic neurons in the substantia nigra parts compacta (SNc). A-syn is a member of a group of intrinsically disordered amyloid proteins that assemble into oligomeric and fibrillar aggregates [4–8].

The exact aetiology of PD is still unknown. The most important risk factor for developing PD is advancing age. Variants in genes like *SNCA, GBA1, PINK1, PRKN, and LRRK2*, contribute to the disease pathology, of which the most important risk factor are the mutations in the *GBA1* gene, found in 10-15% of PD patients [9–15]. *GBA1* is located on chromosome 1 and encodes the lysosomal hydrolase glucocerebrosidase (GCase, or acid-β-glucosidase) that catalyses the cleavage of glucosylceramide (GlcCer) and glucosylsphingosine (GlcSph) into glucose and ceramide, or glucose and sphingosine, respectively [16].

A-syn toxic accumulation has been associated with impairment in several cellular functions, involving mitochondria, proteasomes and lysosome, as well as endoplasmic reticulum (ER) stress, axonal transport deficiencies and, ultimately, impaired dopamine synaptic release [17].

Additionally, a-syn oligomers trigger apoptotic pathways via reactive oxygen species (ROS) production [18,19].

The lysosome plays a critical role in the clearance of a-syn; it is physiologically degraded by both the ubiquitin-proteasome system and the autophagy lysosomal pathway (ALP), using mainly chaperone-mediated autophagy (CMA) and macroautophagy [20–24]. Moreover, lysosomal proteins which are crucial for both macroautophagy and CMA and have been found to be defective in the post-mortem brains of PD patients [25]. Patients with *GBA1* mutations exhibit decreased lysosomal GCase enzyme activity in multiple brain regions [26]. Impairment of GCase activity leads to lysosomal dysfunction, obstructing the trafficking and processing of a-syn [27,28]. Conversely, lysosomal dysfunction can be induced by a-syn itself [29], mainly by interrupting the transport of lysosomal hydrolases from the ER to the Golgi apparatus and the lysosome in neurons [30–32]. Consequently, overexpression of a-syn reduces both GCase activity and protein levels, resulting in a reciprocal biochemical connection between a-synuclein and GCase [33–36].

A-syn accumulation can induce ER stress by disrupting protein trafficking and directly interacting with ER membranes. In addition, ER stress exacerbates a-syn toxicity, forming a feedback loop that contributes to neurodegeneration in synucleinopathies [37–39].

In addition to this, a-syn aberrant accumulation has been linked to increased oxidative stress, a cellular status characterized by an accumulation of ROS. The latter is defined by an imbalance in ROS production and the ability of the cell to increase its antioxidant response, causing cellular toxicity at multiple levels, ranging from altered mitochondrial function to membrane damage.

Microgravity, the condition that is naturally experienced by astronauts in space, has been shown to induce oxidative stress by increasing ROS productions and lipid peroxidation. Space agencies such as the National Aeronautics and Space Administration (NASA) and the European Space Agency (ESA) conduct experiments aboard space stations like the International Space Station (ISS) to explore phenomena not observable under terrestrial gravity (1g). Microgravity profoundly affects biological systems, from single cells to entire organisms and cellular adjustment in these conditions can help the exploration of diverse pathophysiological processes [40–48].

Simulated microgravity (s-μg) environment also impairs cellular stress responses, including the unfolded protein response and protein turnover, which may, in turn, lead to the accumulation of misfolded protein.

These findings imply a potential induction of senescence across various cellular models, including cancer cell lines, stem cells, and differentiated cells [49–52]. While further investigation is necessary to fully understand the implications of microgravity on senescence, several researchers propose that exposure to microgravity triggers cellular aging, possibly through heightened oxidative stress resulting from mitochondrial dysfunction [44,53–58]. This process may initiate apoptotic pathways and alter cellular proliferation, differentiation, and growth [51,59].

In the present study, a systematic characterization of PD features under s-μg conditions using a random positioning machine (RPM) was performed, focusing on a-syn alterations, including the formation of aggregates, lysosomal physiology, and oxidative stress. We employed two neuronal cell line models, the SH-SY5Y and a-syn mutant 3K-SNCA line. Whilst the first cell line is widely used as a model for neurodegenerative disorders [60,61], the mutant line is designed to accelerate a-syn aggregation. Specifically, the familial PD-linked mutation E46K, which changes the a-syn repeat motif KTKEGV to KTKKGV in repeat 4, was amplified by inserting analogous mutations into the immediately adjacent KTKEGV repeat motifs (E35K+E46K+E61K), stabilizing membrane-associated helical a-syn, hence promoting the protein oligomerization and eventual aggregation [62,63].

Our study shows that s-μg conditions result in a-syn alterations and oxidative stress in mutated and wild type cells, indicating that s-μg exposure leads to increased a-syn accumulation and aggregation, independently of the presence of a-syn mutations.

## Methods

### Cell culture

Human neuroblastoma cells SH-SY5Y (ATCC number: CRL-2266) and 3K-SNCA cells were cultured in DMEM/F-12, GlutaMAX™ medium (10565018; Gibco) supplemented with 10% fetal bovine serum (FBS) and 1x Penicillin/Streptomycin (Pen/Strep) at 37° C and 5% CO_2_ in T-75 flasks.

### Microgravity

The Random Positioning Machine (RPM) was used as a microgravity simulator, while, although it cannot be directly compared to the real condition of microgravity, it still serves as a valuable tool for conducting preliminary tests [64]. It is composed by two frames that rotate independently. The samples, which are placed close to the centre in the inner frame, are subjected to a mean gravity force close to zero g, due to the continuous re-orientations and random movements of the axes.

The flasks containing the cells were carefully filled up with complete medium to avoid the shear forces and to allow the cells to maintain a state of free-fall motion. For the experiment, the Random Walk mode was used with a velocity of 60 deg/sec. Cells were left in the machine for different time points (24 hours and 48 hours) and compared with the 1g control group placed in the static bar of the RPM.

### Bradford Assay

Bradford assay was used as a method to quantify the concentration of proteins in samples using bovine serum albumin as a standard according to manufacturer instruction (B6916; Sigma Aldrich).

### Western Blot Analysis

Cells pellets were lysed, and proteins were separated as previously described [65]. The primary antibody were added at the following concentrations: rabbit anti a-syn 1:1000 (ab138501; abcam); mouse anti-GBA1 1:500 (AP1140; Calbio Biochem); mouse anti β-Actin 1:10000 (ab6276; abcam); mouse anti LAMP1 1:500 (611042; BD Trasduction), mouse anti p-Synuclein (Ser129) 1:1000 (ab51253; abcam), rabbit anti Catalase 1:1000 (ab209211; abcam), and mouse anti ATF6 1:500 (ab227830; abcam). The secondary antibody was added at a 1:2000 final concentration: Goat anti-mouse HRP (P0447; Dako) and goat anti-rabbit HRP (P0448; Dako). The membrane was imaged prior incubation with ECL reagent using the ChemiDoc Imaging System.

### Immunostaining and Confocal

Immunostaining was performed as previously described [60,66]. The primary antibodies were added at the following concentrations: mouse anti a-syn antibody 1:750 (SIG-39730; BioLegend) and rabbit anti-LAMP1 1:2000 (ab278043; abcam). The secondary antibodies were added at the following concentrations: 1:2500 Goat anti-Rabbit IgG (H+L) Highly Cross-Adsorbed Secondary Antibody, Alexa Fluor™ Plus 647 (A32733; Invitrogen) and 1:2500 Goat anti-Mouse IgG (H+L) Highly Cross-Adsorbed Secondary Antibody, Alexa Fluor™ Plus 488 (A32723; Invitrogen). 1ng/ml of Hoechst solution (62249; Thermo Scientific) was used to visualize the nuclei. The slides were imaged using the Nikon C2 Confocal Microscope.

### ROS quantification

ROS levels were quantified using Invitrogen™ carboxy-H2DCFDA (C400; Invitrogen) as a fluorescent indicator with a final concentration of 20μM. The experiment was performed on live cells which were prepared by adding 0.5μl of protease inhibitor and anti-phosphatase 1 and 2 to the cell pellets. 1μl of pellet was diluted in 1ml of PBS. H_2_O_2_ was used to prepare a standard curve for different ROS concentrations, using PBS as a control for 0μM ROS. 100 μl of diluted samples and standards were added to a black 96-well plate with 92μl PBS and 8μl carboxy-H2DCFDA probe. After a 1hour incubation in the dark at 37°C with 5% CO_2_, using a filter-based reader (fluorescence), samples were analysed with an excitation of 488nm and emission of 530nm. Final values were normalised for the total amount of protein, quantified using the Bradford assay.

### MDA quantification

The Cayman TBARS assay kit (10009055; Cayman) was performed to measure Malondialdehyde (MDA) levels. In this kit, MDA and Thiobarbituric Acid (TBA) react under high temperature (90-100°C) and acidic conditions, and their product is measured colorimetrically at 530-540nm. MDA standard was diluted to create a stock solution of 125μM, and a standard curve was prepared with varying MDA concentrations. Samples and standards were added to Eppendorf® tubes, along with the SDS solution and the Color reagent (composed of Thiobarbituric Acid, Acetic Acid, and NaOH) according to the manufacturer instructions. The mixture was heated for 1 hour at 100°C, cooled on ice for 10 minutes, centrifuged for 10 minutes at 1,600 x g at 4°C, and then loaded in a clear plate for absorbance reading at 530-540 nm. The standard curve was plotted, and unknown values were interpolated after subtracting the blank. Final values were normalised for the total amount of proteins, quantified using the Bradford assay.

### Total Antioxidant Capacity (TAC) quantification

Cayman’s Antioxidant Assay (709001, Cayman) is a versatile method for assessing the total antioxidant capacity (TAC) of various biological samples. This assay evaluates the combined antioxidant activities of a range of constituents, such as vitamins, proteins, lipids, glutathione, and uric acid.

All the reagents and samples were prepared and diluted according to manufacturer instructions. Trolox standard wells received 10μl of Trolox standard (from a concentration of 0µM to 0.33 µM as suggested in the assay protocol), 10μl of metmyoglobin, and 150μl of chromogen per well. Similarly, sample wells were prepared with 10μl of sample, 10μl of metmyoglobin, and 150μl of chromogen. The reactions were initiated by adding 40μl of hydrogen peroxide working solution to all the wells. It is crucial to add the hydrogen peroxide quickly (within one minute is recommended when using a multichannel) to ensure uniform reaction initiation across wells. The plate was then covered and incubated on a shaker for five minutes at room temperature to allow for the completion of the reactions. After incubation, the plate cover was removed, and the absorbance was measured at 405 nm using a plate reader. The standard curve was plotted, and unknown values were interpolated after subtracting the blank, values were then multiplied for the dilution factor. Final values were normalised for the total amount of proteins, quantified using the Bradford assay.

### DATA analysis

Statistical comparisons and analysis were performed using GraphPad Prism 9.1 software. The Shapiro-Wilk test was first used to determine whether the collected data was normally distributed. In non-normally distributed data, Kruskal-Wallis test and Dunn’s post-hoc analysis were used to conduct multiple comparisons. The differences between three or more groups of normally distributed samples were compared using the ANOVA test. Each experiment was conducted three times. Each condition was carried out in triplicate in every experiment conducted. Results are displayed as mean ± standard error of the mean. Results were considered statistically significant if their p-value was less than 0.05.

## Results

### A-syn pathogenic species are modulated by s-**μ**g exposure

To study whether s-μg exposure influences a-syn protein behaviour, we quantified various species of the protein. Initially, we questioned whether the total level of a-syn is affected by prolonged exposure to s-μg. Thus, two different time points (24 and 48 hours) were assessed to monitor the changes in a-syn levels. We then studied whether a change in total levels of a-syn would translate in the different subspecies of the protein. We quantified aggregated forms of a-syn which are known to impair lysosomal function, induce ER stress, and promote an aberrant UPR [26,67,68].

The initial investigation on total a-syn levels, using a western blot approach, did not reveal any significant difference between cells exposed s-μg and those cultured in 1g, independently of the SNCA mutation (Fig. 1 a-d). These findings suggest that the overall quantity of a-syn in cell cultures in s-μg is not affected. However, confocal microscopy analysis of 3K-SNCA cells exposed to s-μg revealed a significant increase in a-syn aggregates after just 24 hours of exposure (P=0.0007), with a further marked increase following 48 hours of incubation in the s-μg gravity environment (P=0.000006) (Fig. 2 a, b). To ensure that the observed effects were not due to a combined influence of the 3K mutation and s-μg, we exposed the SH-SY5Y control cell line to s-μg. This experiment revealed a significant increase in a-syn aggregates at both 24 hours (P=0.0076) and 48 hours of exposure (P=0.0364) (Fig. 2 c, d). The detection of aberrant a-syn accumulation in the SH-SY5Y control line confirms that the observed increase was indeed driven by s-μg and not an exacerbation of an existing defect, such as the 3K-SNCA mutation. Following the identification of increased presence of a-syn aggregates, we questioned whether these findings were driven by a constitutive increase in the a-syn monomeric fraction, acting as a potential additional protein source and thereby promoting the formation of aggregated forms. To investigate the levels of a-syn monomers, we quantified the remaining a-syn signal that was not identified as an “aggregate” throughout the confocal microscopy screening. This assessment revealed a significant depletion of a-syn monomers in both 3K-SNCA cells (P=0.0145; P=0.0025) and SH-SY5Y cells (P=0.039; P=0.015) following s-μg exposure at both time points analyzed (Fig. 2 e, f).

**Figure 1.**
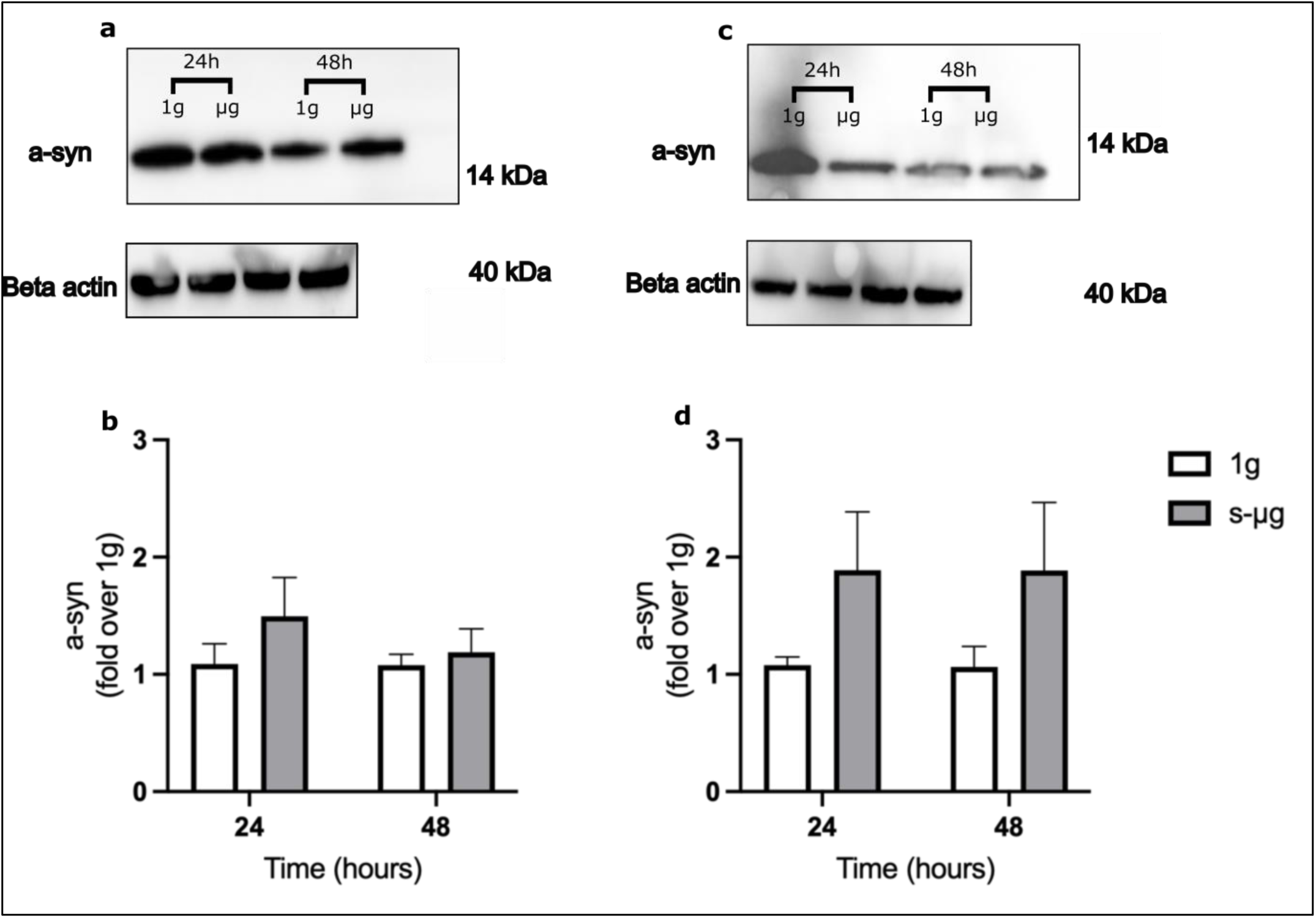
Total a-syn quantification in SH-SY5Y and 3K-SNCA cells exposed to s-mg conditions at different time points. a) Representative WB membrane image of a-syn and control protein beta-actin in SH-SY5Y cells. b) Quantification of a-syn levels in 1g and s-mg exposed SH-SY5Y cells. c) Representative WB membrane image of a-syn and control protein beta-actin in 3K-SNCA cells. d) Quantification of a-syn levels in 1g and s-mg exposed SH-SY5Y cells. Multiple unpaired t-tests were used to compare differences. Data are presented as mean±SEM of 3 independent experiments.

**Figure 2.**
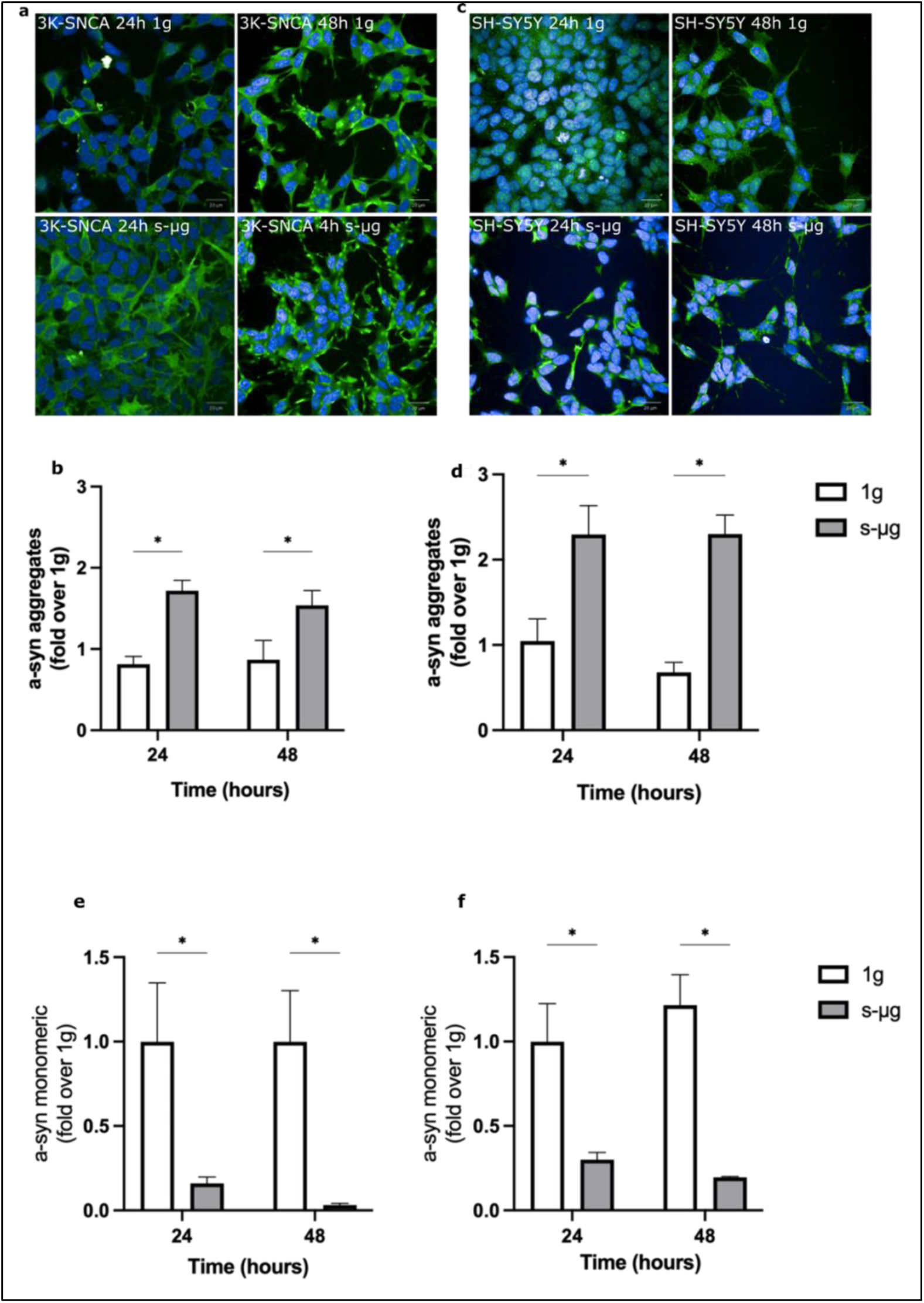
Quantification of a-syn aggregates via confocal microscopy in SH-SY5Y cells and 3K-SNCA cells. a-syn (green), Nuclei (blue). **a)** Representative confocal membrane images of a-syn aggregates in 3K-SNCA cells cultured in 1g and s-μg. **b)** Quantification of a-syn aggregates in 1g and s-μg exposed 3K-SNCA cells cultured in 1g and s-μg. **c)** Representative confocal images of a-syn aggregates in SH-SY5Y cells. **d)** Quantification of a-syn aggregates levels in 1g and s-μg exposed SH-SY5Y cells. **e)** Quantification of non-aggregated a-syn in 1g and s-μg exposed 3K-SNCA cells cultured in 1g and s-μg. **f)** Quantification of non-aggregated a-syn in 1g and s-μg exposed SH-SY5Y cells cultured in 1g and s-μg. Multiple unpaired t-tests were used to compare differences. Data are presented as mean±SEM of 3 independent experiments. \**P*<0.05

To understand whether the observed increase in aggregated a-syn following s-μg exposure was accompanied an initial shift in pro-aggregation forms of the protein, we quantified the levels phosphorylated form on Serine-129 of a-syn (pS219). This phosphorylated form is a well-established marker of the aggregated state, as approximately 90% of a-synuclein in Lewy Bodies from PD brains contains pS129. Moreover, pS129 has been shown to accelerate the rate of a-syn fibrillation in vitro, suggesting that this specific phosphorylation event is closely linked to pathological aggregation.[69–71].

Initially, we conducted a comparative analysis between 1g SH-SY5Y cells and 1g 3K-SNCA cells to ascertain variations in pS219 presence. Our findings revealed a notably higher abundance of pS219 in 3K-SNCA cells compared to the wild-type model, substantiating and aligning with previous reports on the literature (data not shown) [72]. Levels of pS129 were significantly elevated in both cell types following exposure to s-μg, with this increase becoming more pronounced over the two time points analyzed (Fig. 3 a-d). These findings suggest that under s-μg conditions a-syn shifts from a non-aggregated status to phosphorylated and aggregated forms.

**Figure 3.**
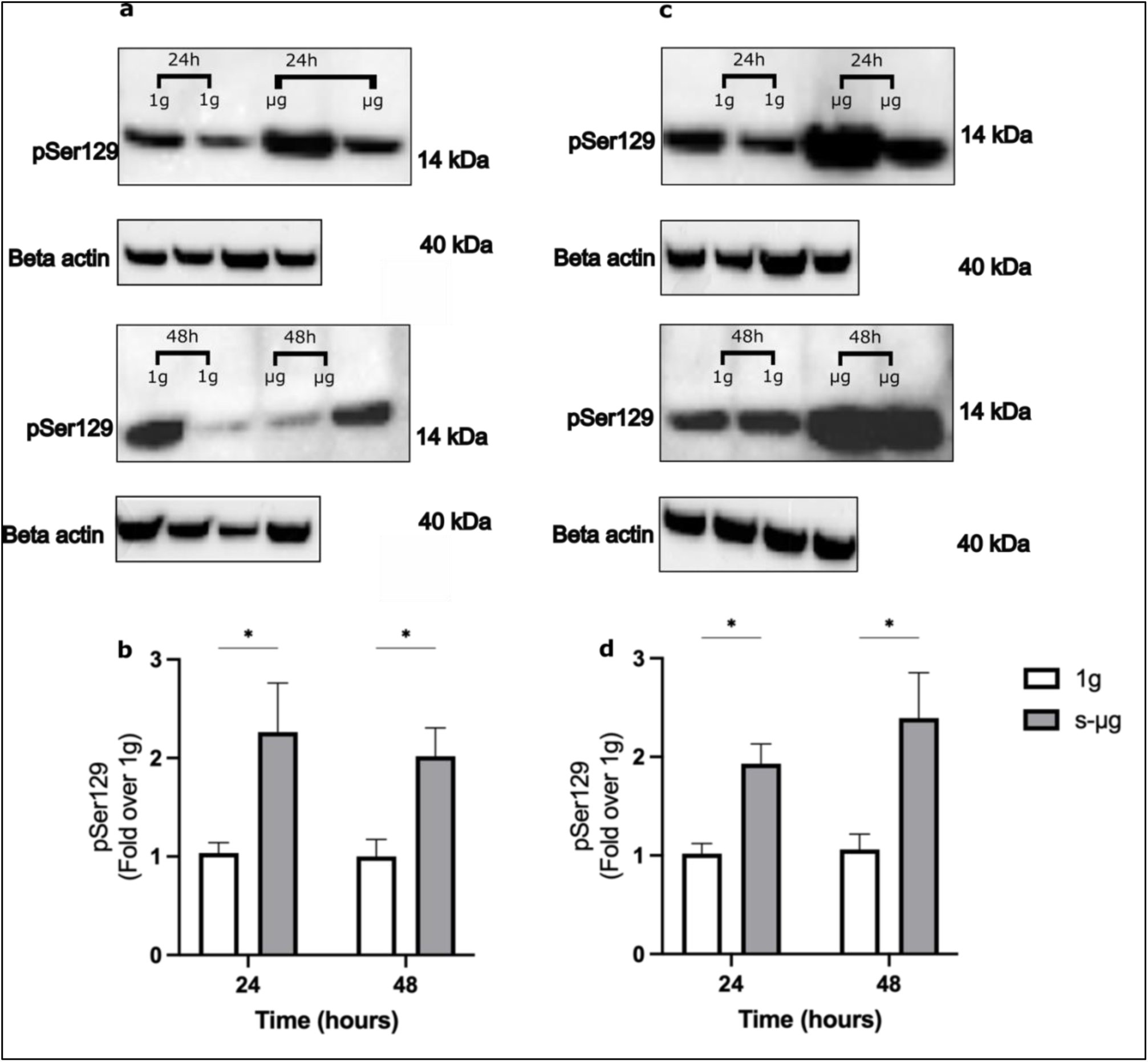
phosphorylated Serine 129 a-syn (pSer129) quantification in SH-SY5Y and 3K-SNCA cells exposed to s-μg at different time points. **a)** Representative WB membrane image of pSer129 and control protein beta-actin in SH-SY5Y cells. **b)** Quantification of pSer129 levels in 1g and s-μg exposed SH-SY5Y cells. **c)** Representative WB membrane image of pSer129 and control protein beta-actin in 3K-SNCA cells. **d)** Quantification of pSer129 levels in 1g and s-μg exposed SH-SY5Y cells. Multiple unpaired t-tests were used to compare differences. Data are presented as mean±SEM of 3 independent experiments. \**P*<0.05

### GCase is not increased in s-μg

Given that s-μg may impact cellular processes, the abnormal behavior of a-syn observed under these conditions might be due to disrupted lysosomal function, leading to impaired clearance and increased aggregation of the protein. Thus, we hypothesized that this could be mechanistically related to lysosomal dysfunction. To investigate this, we quantified the levels of GCase upon exposition of the cells to s-μg conditions using the RPM. GCase was selected as it is an important lysosomal enzyme in PD and a-syn turnover. Our analysis revealed that GCase levels were not significantly altered at 24 or 48 hours (Fig. 4 a-d). We also measured Lysosomal-Associated Membrane Protein 1 (LAMP1) levels as a reflection of lysosomal mass. Quantification of LAMP1 punctae using confocal microscopy revealed a significant increase in LAMP1-positive organelles over time under s-μg conditions. Specifically, SH-SY5Y cells exhibited a more than 2-fold increase at both 24 hours (P=0.039) and 48 hours (P=0.025) of s-μg exposure. The PD model line 3K-SNCA presented more variable levels of LAMP1, with a significant increase only present in exposure to s-μg at 48 hours (*P=*0.003). Although requiring further investigation, these results suggest that LAMP1 increase may not be followed by a simultaneous increase in lysosomal hydrolases like GBA1, potentially resulting in an impaired lysosomal maturation in s-μg.

**Figure 4.**
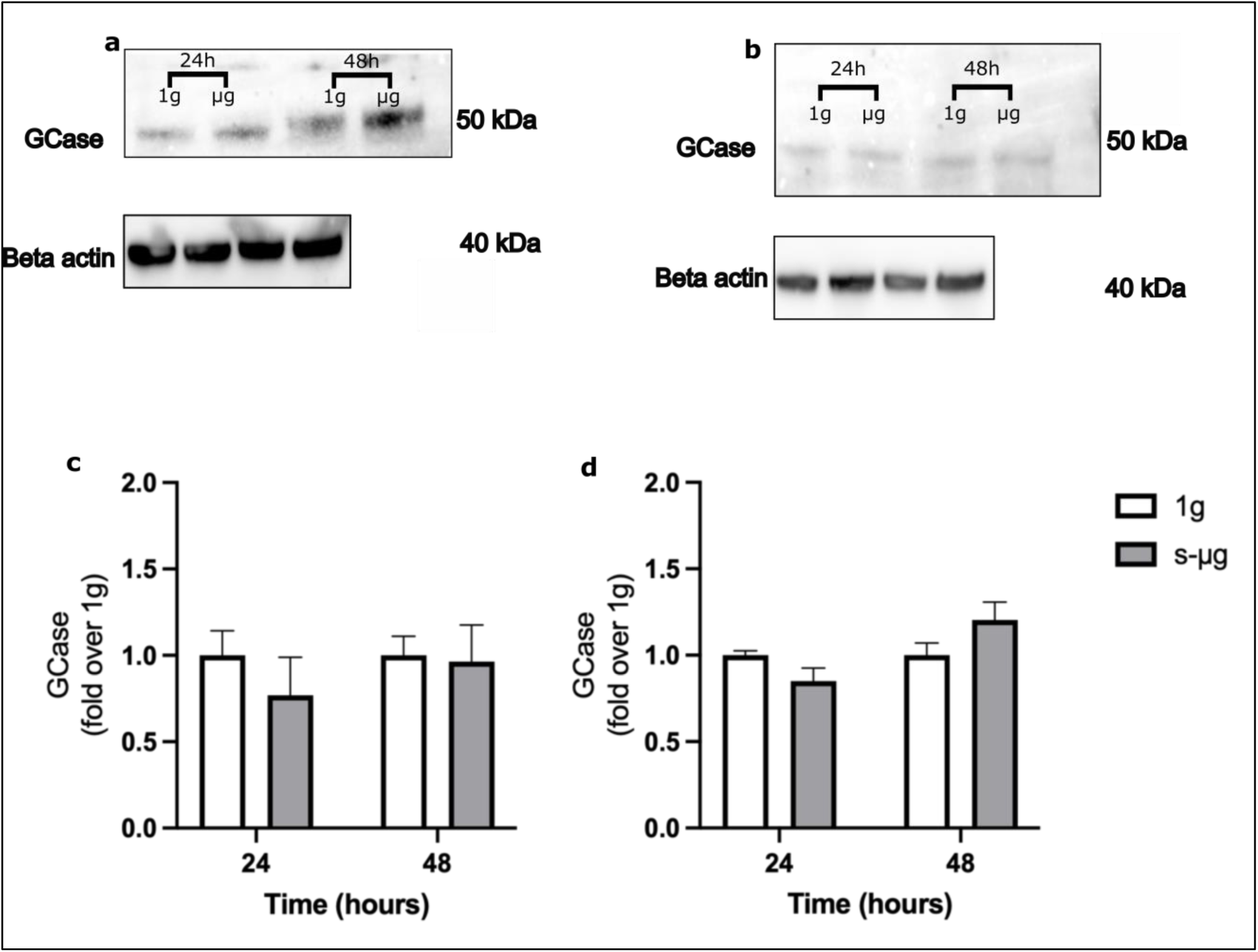
GCase WB quantification in SH-SY5Y and 3K-SNCA cells exposed to s-μg conditions at different time points. **a)** Representative WB membrane image of GCase and control protein beta-actin in SH-SY5Y cells. **b)** Quantification of GCase levels in 1g and s-μg exposed SH-SY5Y cells. **c)** Representative WB membrane image of GCase and control protein beta-actin in 3K-SNCA cells. **d)** Quantification of GCase levels in 1g and s-μg exposed SH-SY5Y cells. Multiple unpaired t-tests were used to compare differences. Data are presented as mean±SEM of 3 independent experiments.

### ROS levels are increased in s-μg conditions

When subjected to the s-μg induced by the random motion of the machine, both the 3K-SNCA overexpressing cells and the control SH-SY5Y cells exhibited an elevated quantity of both monomeric and aggregated states. An increase in oxidative stress is seen in the brain in several neurodegenerative diseases including PD and has been noted in microgravity cell models previously[56,57]. To investigate oxidative stress in our a-syn models, we quantified ROS and lipid peroxidation levels in SH-SY5Y and 3K-SNCA cells exposed to s-μg conditions at 24 hours and 48 hours.

SH-SY5Y cells showed a significant 3-fold (*P=*0.019) and 2-fold increase (*P=*0.02) in ROS at 24 hours and 48 hours respectively. Similarly, 3K-SNCA cells showed a significant increase of ROS cellular content at both time points analysed (*P=*0.015; *P=*0.031) (Fig. 6 a, b).

**Figure 5.**
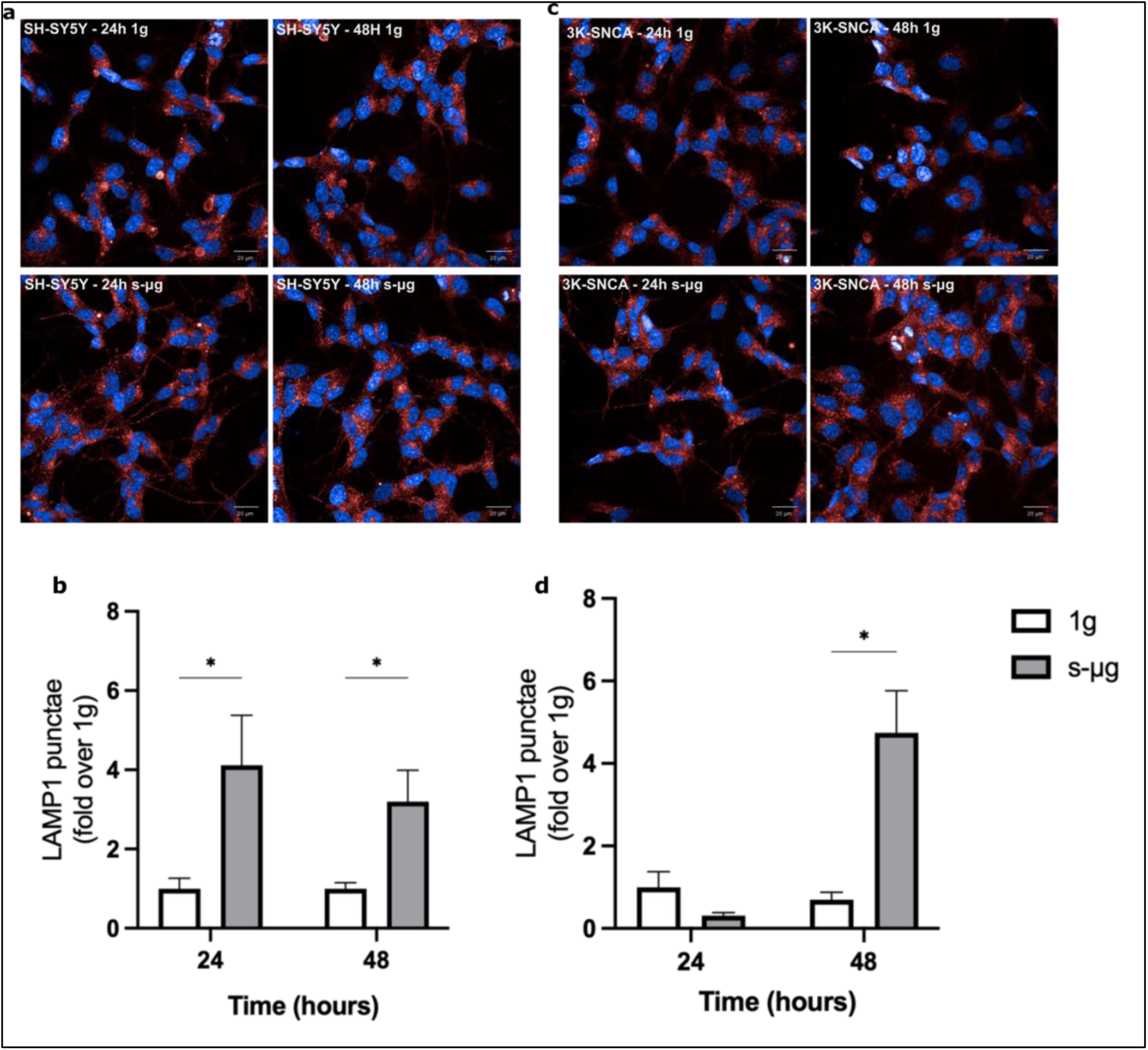
Quantification of LAMP1 punctae via confocal microscopy in SH-SY5Y cells and 3K-SNCA cells. LAMP1 (red), Nuclei (blue). **a)** Representative confocal membrane images of LAMP1 punctae in SH-SY5Y cells cultured in 1g and s-μg. **b)** Quantification of LAMP1 punctae in 1g and s-μg exposed SH-SY5Y cells. **c)** Representative confocal images of LAMP1 punctae in 3K-SNCA cells. **d)** Quantification of LAMP1 punctae levels in 1g and s-μg exposed 3K-SNCA cells. Multiple unpaired t-tests were used to compare differences. Data are presented as mean±SEM of 3 independent experiments. \**P*<0.05

**Figure 6.**
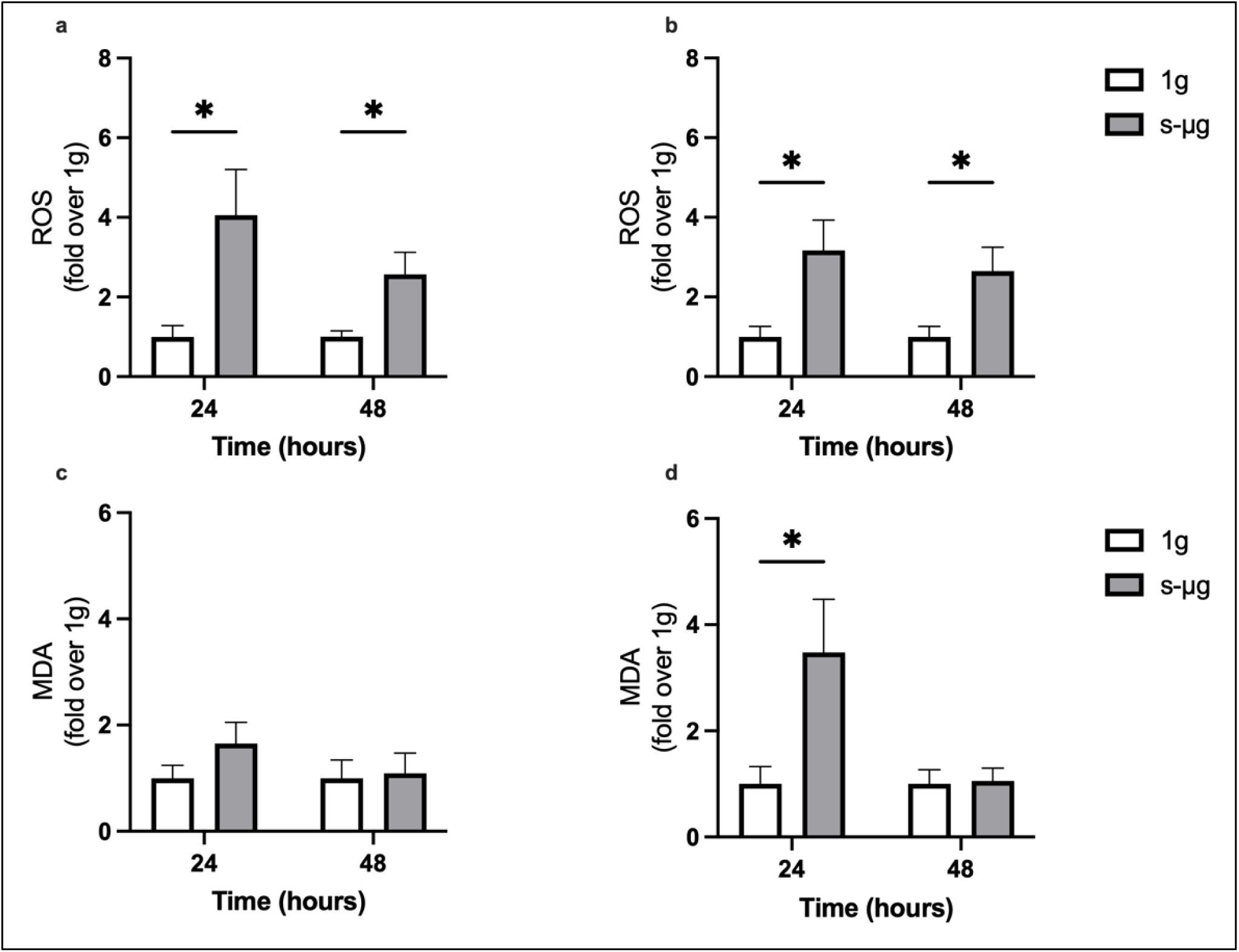
Investigation on oxidative stress marker in SH-SY5Y cells and 3K-SNCA cells exposed to s-μg. **a)** Quantification of ROS levels in SH-SY5Y cells cultured in 1g and s-μg. **b)** Quantification of ROS levels in 3K-SNCA cells cultured in 1g and s-μg. **c)** Quantification of MDA level in 1g and s-μg exposed SH-SY5Y cells. **d)** Quantification of MDA level in 1g and s-μg exposed 3K-SNCA cells. Multiple unpaired t-tests were used to compare differences. Data are presented as mean±SEM of 3 independent experiments. \**P*<0.05

We also measured MDA levels as an indicator of lipid peroxidation. MDA is generated by the peroxidation of membrane polyunsaturated fatty acids [73] as well as produced in the process of prostaglandin synthesis [74]. As s-μg conditions are a source of oxidative stress, s-μg also increases MDA levels [75]. We exposed 3K-SNCA and SH-SY5Y cells to s-μg conditions for 24 and 48 hours, using standard gravity exposed-cells as a control. There was no significant change in MDA levels in the SH-SY5Y cells at either time point. In the 3K-SNCA, however, there was a significant 3-fold increase (*P=*0.025) at 24 but not 48 hours (Fig. 6 c, d). These data are consistent with previous studies showing that microgravity results in an increased presence of oxidative stress markers within the cells. The failure of MDA levels to rise in the SH-SY5Y cells and the reversal of increased levels between 24 and 48 hours in the 3K-SNCA cells suggests that compensatory mechanisms are modulated in response to the increased ROS. For this, we investigate the levels of Catalase, responsible of H_2_O_2_ hydrolysis, along with the TAC. However, we did not find any change in both Catalase (Fig. S1 a-d) and TAC (Fig. S2 a-b) in either cell at any time point suggesting a potential failure of antioxidant response despite the increased presence of ROS following s-μg exposure.

## Discussion

Microgravity research is increasingly recognized for its potential across various scientific disciplines. While further investigation is needed to fully understand cellular adaptation to space-like conditions, exploring how cells adjust to microgravity offers substantial promise for revealing diverse pathophysiological processes [44,54,78,79]. In this study, s-μg was employed to simulate accelerated aging, providing insights into the underlying pathophysiological mechanisms of PD. We report that s-μg increases a-syn aggregate levels in both SH-SY5Y and 3K-SNCA cell models. Notably, exposure to s-μg shifts the a-syn ratio towards aggregated form and increased pS129 modification, both favoring the potential for Lewy body-like pathology [56,57].

All these observations could be attributed to s-μg induced cell stress, conceivably leading eventually to a diminished clearance of a-syn, particularly pronounced in 3K-SNCA overexpressing cells. For this reason, it was essential to investigate whether the increase in a-syn relates to compromised protein clearance mechanisms, specifically involving autophagy–lysosome system dysfunction.

We hypothesised that lysosomal biogenesis was upregulated in s-μg as compensation for accumulated a-syn aggregates. To evaluate this, we quantified both GCase lysosomal hydrolase as well as LAMP1 in both cell types. Our analyses revealed that GCase levels remains unchanged, whilst LAMP1 increases over time in both cell types. Several studies have demonstrated that GCase homeostasis is affected by a-syn accumulation [32] by causing ER fragmentation and compromising ER folding capacity and facilitating the aggregation of lysosomal hydrolases in the ER. The failure for GCase levels to rise in our study may reflect this effect of a-syn, even against a compensatory attempt to increase the endo-lysosomal content [30,31,46]. Accumulation of both a-syn and immature lysosome hydrolases in the ER leads to activation of ER stress, thus evoking UPR [74,76] and resulting in loss of ER homeostasis. This potentially culminates in ER dysfunction (changes in ER calcium concentration or dysregulation in the redox potential of the ER) [77] or indirectly lysosomal catabolic malfunction or mitochondrial abnormalities, which in turn could results in harmful oxidative damage to the cells, and further a-syn aggregation [78,79].

In addition to this, ROS levels were increased in both cell types under s-μg conditions. Both s-μg and a-syn aggregates accumulation are known to increase oxidative stress [19,80–82]. Additionally, a-syn has been reported to suppress catalase activity [83]. These results suggests that cells exposed to s-μg fail to activate protective mechanisms against rising levels of ROS, which may contribute to an endo-lysosomal impairment, promoting GCase lysosomal hydrolase aberrant homeostasis, potentially further facilitating a-synuclein aggregation.

Our findings describe a senescence model of PD cell pathology induced by microgravity, with misfolded a-syn accumulation and oxidative stress. While the characteristics of the cell models used in our study offer valuable insights for specific experimental contexts, they do not fully replicate the complex cellular dynamics observed in PD. In contrast, differentiated cells, which more closely resemble the specialized neuronal populations, would provide a more accurate representation of PD pathological processes. These cells would better capture the scenario of neuronal dysfunction, including the impact of a-syn aggregation and oxidative stress on cellular health and will be used in future studies.

## Conclusions

We report the first studies on the effect of s-μg on a-syn and show that it induces and enhances a-syn accumulation in neuronal-type cells, accompanied by an increase in oxidative stress. These findings provide valuable insights into cellular responses to s-μg, providing a basis for the usage of s-μg and space-based experiments to model complex neurodegenerative disorders and ageing features.

## Acknowledgments

This work was funded by e.INS-Ecosystem of Innovation for Next Generation Sardinia (Grant ECS 00000038) funded by the Italian Ministry for Research and Education (MUR) under the National Recovery and Resilience Plan (NRRP) MISSION 4COMPONENT 2, “From Research to Business” INVESTMENT 1.5, “Creation and Strengthening of Ecosystems of Innovation”, and construction of “Territorial R&D Leaders”.

## Authors contributions

VL and GU contributed equally to this research. VL and GU conceived the study, designed and executed the analysis plan with supervision from GC, AP, and AHVS. GU, VL, AM, MAEF, and ND performed tissue culture work, immunostaining and data collection. VL and GU performed image analysis and conducted statistical analysis. AP and AHVS supported data interpretation. VL, GU, SLDP and AHVS wrote, designed, drafted and edited the manuscript with feedback and input from all authors. All authors read and approved the final manuscript.

## Conflict of intertest

Nothing to report.

## Data availability

The datasets in this study are not publicly available at the moment. They will be publicly available once the manuscript is accepted for publication.

## Figures

**Figure S1.**
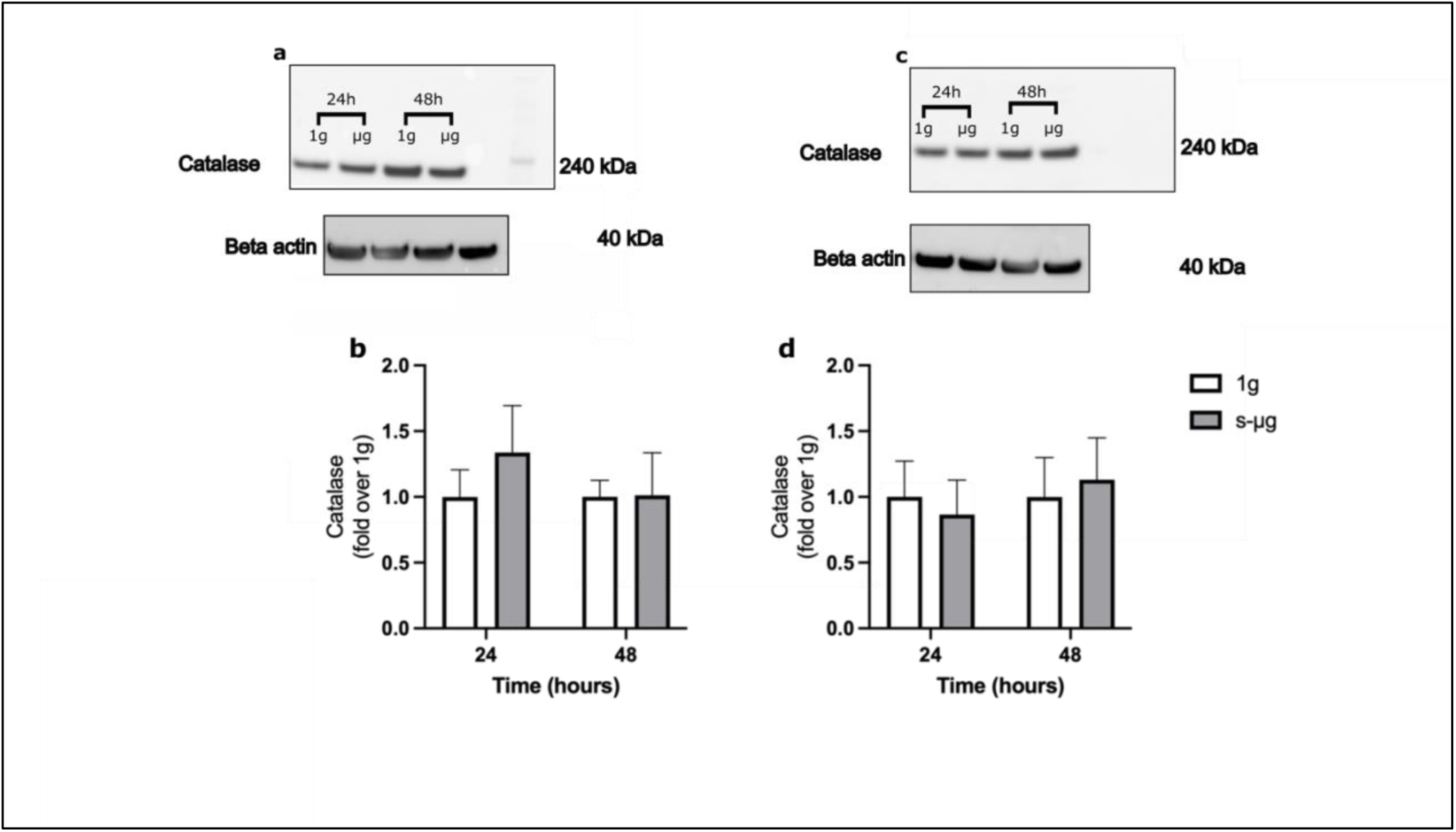
Catalase WB quantification in SH-SY5Y and 3K-SNCA cells exposed to s-μg at different time points. a) Representative WB membrane image of Catalase and control protein beta-actin in SH-SY5Y cells. b) Quantification of Catalase levels in 1g and s-μg exposed SH-SY5Y cells. c) Representative WB membrane image of Catalase and control protein beta-actin in 3K-SNCA cells. d) Quantification of Catalase levels in 1g and s-μg exposed SH-SY5Y cells. Multiple unpaired t-tests were used to compare differences. Data are presented as mean±SEM of 3 independent experiments.

**Figure S2.**
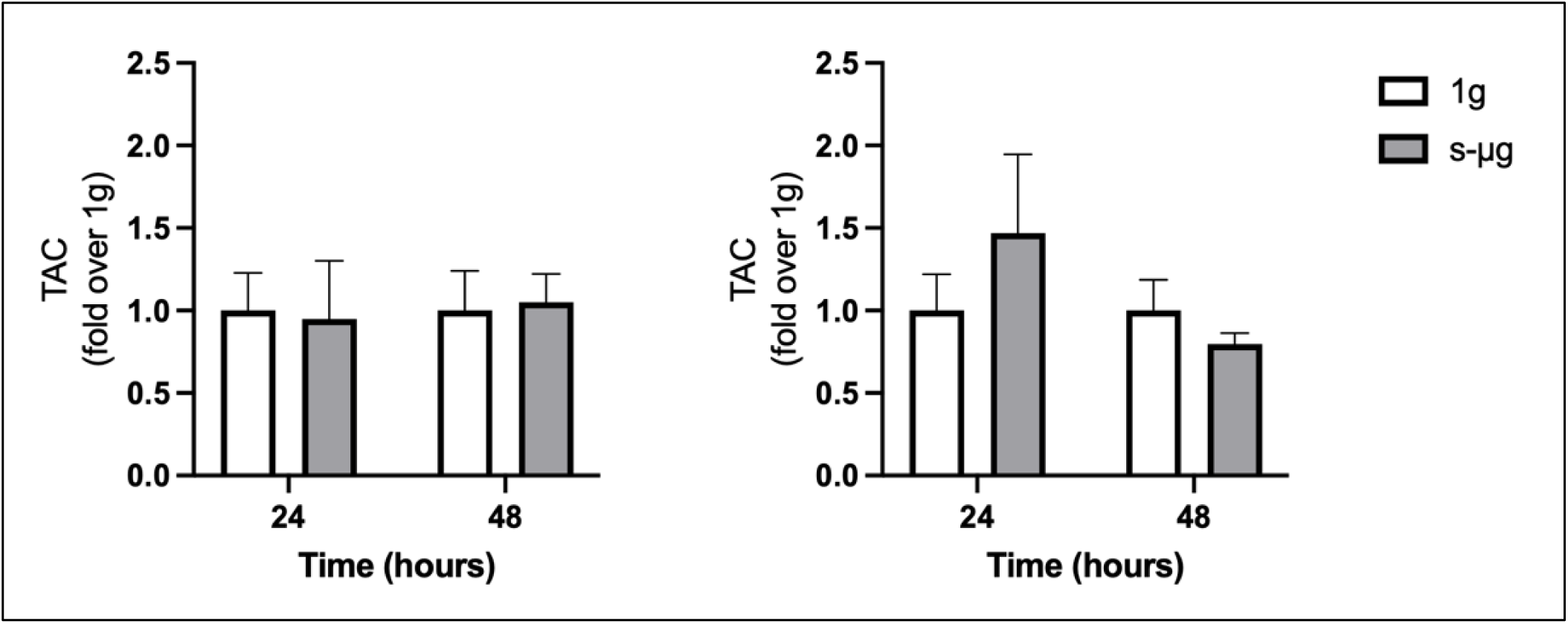
Investigation on total antioxidant capacity (TAC) in SH-SY5Y cells and 3K-SNCA cells exposed to s-μg. a) Quantification of TAC level in SH-SY5Y cells cultured in 1g and s-μg. b) Quantification of TAC level in 3K-SNCA cells cultured in 1g and s-μg. c) Quantification of TAC level in 1g and s-μg exposed SH-SY5Y cells. d) Quantification of TAC level in 1g and s-μg exposed 3K-SNCA cells. Multiple unpaired t-tests were used to compare differences. Data are presented as mean±SEM of 3 independent experiments.

## Bibliography

1. Parkinson Disease Available online: https://www.who.int/news-room/fact-sheets/detail/parkinson-disease (accessed on 19 January 2023).

2. Postuma, R.B.; Berg, D.; Stern, M.; Poewe, W.; Olanow, C.W.; Oertel, W.; Obeso, J.; Marek, K.; Litvan, I.; Lang, A.E.;, et al. MDS Clinical Diagnostic Criteria for Parkinson’s Disease. Movement Disorders 2015, 30.

3. Schapira, A.H.V.; Chaudhuri, K.R.; Jenner, P. Non-Motor Features of Parkinson Disease. Nat Rev Neurosci 2017, 18.

4. Poewe, W.; Seppi, K.; Tanner, C.M.; Halliday, G.M.; Brundin, P.; Volkmann, J.; Schrag, A.E.; Lang, A.E. Parkinson Disease. Nature Reviews Disease Primers 2017 3:1 2017, 3, 1–21, doi:10.1038/nrdp.2017.13.

5. Balestrino, R.; Schapira, A.H.V. Parkinson Disease. Eur J Neurol 2020, 27, 27–42, doi:10.1111/ENE.14108.

6. Spillantini, M.G.; Schmidt, M.L.; Lee, V.M.Y.; Trojanowski, J.Q.; Jakes, R.; Goedert, M. α-Synuclein in Lewy Bodies [8]. Nature 1997, 388.

7. Shahmoradian, S.H.; Lewis, A.J.; Genoud, C.; Hench, J.; Moors, T.E.; Navarro, P.P.; Castaño-Díez, D.; Schweighauser, G.; Graff-Meyer, A.; Goldie, K.N.;, et al. Lewy Pathology in Parkinson’s Disease Consists of Crowded Organelles and Lipid Membranes. Nat Neurosci 2019, 22, doi:10.1038/s41593-019-0423-2.

8. Mehra, S.; Sahay, S.; Maji, S.K. α-Synuclein Misfolding and Aggregation: Implications in Parkinson’s Disease Pathogenesis. Biochim Biophys Acta Proteins Proteom 2019, 1867.

9. Polymeropoulos, M.H.; Lavedan, C.; Leroy, E.; Ide, S.E.; Dehejia, A.; Dutra, A.; Pike, B.; Root, H.; Rubenstein, J.; Boyer, R.;, et al. Mutation in the Alpha-Synuclein Gene Identified in Families with Parkinson’s Disease. Science 1997, 276, 2045–2047, doi:10.1126/SCIENCE.276.5321.2045.

10. Singleton, A.B.; Farrer, M.; Johnson, J.; Singleton, A.; Hague, S.; Kachergus, J.; Hulihan, M.; Peuralinna, T.; Dutra, A.; Nussbaum, R.;, et al. Alpha-Synuclein Locus Triplication Causes Parkinson’s Disease. Science 2003, 302, 841, doi:10.1126/SCIENCE.1090278.

11. Sidransky, E.; Nalls, M.A.; Aasly, J.O.; Aharon-Peretz, J.; Annesi, G.; Barbosa, E.R.; Bar-Shira, A.; Berg, D.; Bras, J.; Brice, A.;, et al. Multicenter Analysis of Glucocerebrosidase Mutations in Parkinson’s Disease. N Engl J Med 2009, 361, 1651–1661, doi:10.1056/NEJMOA0901281.

12. Tan, E.K.; Yew, K.; Chua, E.; Puvan, K.; Shen, H.; Lee, E.; Puong, K.Y.; Zhao, Y.; Pavanni, R.; Wong, M.C.;, et al. PINK1 Mutations in Sporadic Early-Onset Parkinson’s Disease. Movement Disorders 2006, 21, 789–793, doi:10.1002/MDS.20810.

13. Kitada, T.; Asakawa, S.; Hattori, N.; Matsumine, H.; Yamamura, Y.; Minoshima, S.; Yokochi, M.; Mizuno, Y.; Shimizu, N. Mutations in the Parkin Gene Cause Autosomal Recessive Juvenile Parkinsonism. Nature 1998 392:6676 1998, 392, 605–608, doi:10.1038/33416.

14. Zimprich, A.; Biskup, S.; Leitner, P.; Lichtner, P.; Farrer, M.; Lincoln, S.; Kachergus, J.; Hulihan, M.; Uitti, R.J.; Calne, D.B.;, et al. Mutations in LRRK2 Cause Autosomal-Dominant Parkinsonism with Pleomorphic Pathology. Neuron 2004, 44, 601–607, doi:10.1016/J.NEURON.2004.11.005.

15. Schapira, A.H.; Jenner, P. Etiology and Pathogenesis of Parkinson’s Disease. Movement Disorders 2011, 26, 1049–1055, doi:10.1002/MDS.23732.

16. Berent, S.L.; Radin, N.S. Mechanism of Activation of Glucocerebrosidase by CO-β-Glucosidase (Glucosidase Activator Protein). Biochimica et Biophysica Acta (BBA)/Lipids and Lipid Metabolism 1981, 664, doi:10.1016/0005-2760(81)90134-X.

17. Bendor, J.T.; Logan, T.P.; Edwards, R.H. The Function of α-Synuclein. Neuron 2013, 79.

18. Du, X.Y.; Xie, X.X.; Liu, R.T. The Role of α-Synuclein Oligomers in Parkinson’s Disease. Int J Mol Sci 2020, 21, 1–17, doi:10.3390/IJMS21228645.

19. Deas, E.; Cremades, N.; Angelova, P.R.; Ludtmann, M.H.R.; Yao, Z.; Chen, S.; Horrocks, M.H.; Banushi, B.; Little, D.; Devine, M.J.;, et al. Alpha-Synuclein Oligomers Interact with Metal Ions to Induce Oxidative Stress and Neuronal Death in Parkinson’s Disease. Antioxid Redox Signal 2016, 24, 376–391, doi:10.1089/ARS.2015.6343.

20. Tanik, S.A.; Schultheiss, C.E.; Volpicelli-Daley, L.A.; Brunden, K.R.; Lee, V.M.Y. Lewy Body-like α-Synuclein Aggregates Resist Degradation and Impair Macroautophagy. Journal of Biological Chemistry 2013, 288, doi:10.1074/jbc.M113.457408.

21. Ebrahimi-Fakhari, D.; Cantuti-Castelvetri, I.; Fan, Z.; Rockenstein, E.; Masliah, E.; Hyman, B.T.; McLean, P.J.; Unni, V.K. Distinct Roles in Vivo for the Ubiquitin-Proteasome System and the Autophagy-Lysosomal Pathway in the Degradation of α-Synuclein. Journal of Neuroscience 2011, 31, doi:10.1523/JNEUROSCI.1560-11.2011.

22. Vogiatzi, T.; Xilouri, M.; Vekrellis, K.; Stefanis, L. Wild Type α-Synuclein Is Degraded by Chaperone-Mediated Autophagy and Macroautophagy in Neuronal Cells. Journal of Biological Chemistry 2008, 283, doi:10.1074/jbc.M801992200.

23. Cuervo, A.M.; Stafanis, L.; Fredenburg, R.; Lansbury, P.T.; Sulzer, D. Impaired Degradation of Mutant α-Synuclein by Chaperone-Mediated Autophagy. Science (1979) 2004, 305, doi:10.1126/science.1101738.

24. Wallings, R.L.; Humble, S.W.; Ward, M.E.; Wade-Martins, R. Lysosomal Dysfunction at the Centre of Parkinson’s Disease and Frontotemporal Dementia/Amyotrophic Lateral Sclerosis. Trends Neurosci 2019, 42.

25. Alvarez-Erviti, L.; Rodriguez-Oroz, M.C.; Cooper, J.M.; Caballero, C.; Ferrer, I.; Obeso, J.A.; Schapira, A.H.V. Chaperone-Mediated Autophagy Markers in Parkinson Disease Brains. Arch Neurol 2010, 67, doi:10.1001/archneurol.2010.198.

26. Gegg, M.E.; Burke, D.; Heales, S.J.R.; Cooper, J.M.; Hardy, J.; Wood, N.W.; Schapira, A.H.V. Glucocerebrosidase Deficiency in Substantia Nigra of Parkinson Disease Brains. Ann Neurol 2012, 72, 455–463, doi:10.1002/ANA.23614.

27. Sidransky, E.; Lopez, G. The Link between the GBA Gene and Parkinsonism. Lancet Neurol 2012, 11, doi:10.1016/S1474-4422(12)70190-4.

28. Kuo, S.H.; Tasset, I.; Cheng, M.M.; Diaz, A.; Pan, M.K.; Lieberman, O.J.; Hutten, S.J.; Alcalay, R.N.; Kim, S.; Ximénez-Embún, P.;, et al. Mutant Glucocerebrosidase Impairs α-Synuclein Degradation by Blockade of Chaperone-Mediated Autophagy. Sci Adv 2022, 8, doi:10.1126/sciadv.abm6393.

29. Mazzulli, J.R.; Zunke, F.; Tsunemi, T.; Toker, N.J.; Jeon, S.; Burbulla, L.F.; Patnaik, S.; Sidransky, E.; Marugan, J.J.; Sue, C.M.;, et al. Activation of β-Glucocerebrosidase Reduces Pathological α-Synuclein and Restores Lysosomal Function in Parkinson’s Patient Midbrain Neurons. Journal of Neuroscience 2016, 36, doi:10.1523/JNEUROSCI.0628-16.2016.

30. García-Sanz, P.; Orgaz, L.; Fuentes, J.M.; Vicario, C.; Moratalla, R. Cholesterol and Multilamellar Bodies: Lysosomal Dysfunction in GBA-Parkinson Disease. Autophagy 2018, 14.

31. García-Sanz, P.; Orgaz, L.; Bueno-Gil, G.; Espadas, I.; Rodríguez-Traver, E.; Kulisevsky, J.; Gutierrez, A.; Dávila, J.C.; González-Polo, R.A.; Fuentes, J.M.;, et al. N370S-GBA1 Mutation Causes Lysosomal Cholesterol Accumulation in Parkinson’s Disease. Movement Disorders 2017, 32, doi:10.1002/mds.27119.

32. Stojkovska, I.; Wani, W.Y.; Zunke, F.; Belur, N.R.; Pavlenko, E.A.; Mwenda, N.; Sharma, K.; Francelle, L.; Mazzulli, J.R. Rescue of α-Synuclein Aggregation in Parkinson’s Patient Neurons by Synergistic Enhancement of ER Proteostasis and Protein Trafficking. Neuron 2022, 110, doi:10.1016/j.neuron.2021.10.032.

33. Smith, L.; Schapira, A.H.V. GBA Variants and Parkinson Disease: Mechanisms and Treatments. Cells 2022, Vol. 11, Page 1261 2022, 11, 1261, doi:10.3390/CELLS11081261.

34. Behl, T.; Kaur, G.; Fratila, O.; Buhas, C.; Judea-Pusta, C.T.; Negrut, N.; Bustea, C.; Bungau, S. Cross-Talks among GBA Mutations, Glucocerebrosidase, and α-Synuclein in GBA-Associated Parkinson’s Disease and Their Targeted Therapeutic Approaches: A Comprehensive Review. Translational Neurodegeneration 2021 10:1 2021, 10, 1–13, doi:10.1186/S40035-020-00226-X.

35. Henderson, M.X.; Cornblath, E.J.; Darwich, A.; Zhang, B.; Brown, H.; Gathagan, R.J.; Sandler, R.M.; Bassett, D.S.; Trojanowski, J.Q.; Lee, V.M.Y. Spread of α-Synuclein Pathology through the Brain Connectome Is Modulated by Selective Vulnerability and Predicted by Network Analysis. Nat Neurosci 2019, 22, doi:10.1038/s41593-019-0457-5.

36. Mazzulli, J.R.; Xu, Y.H.; Sun, Y.; Knight, A.L.; McLean, P.J.; Caldwell, G.A.; Sidransky, E.; Grabowski, G.A.; Krainc, D. Gaucher Disease Glucocerebrosidase and α-Synuclein Form a Bidirectional Pathogenic Loop in Synucleinopathies. Cell 2011, 146, doi:10.1016/j.cell.2011.06.001.

37. Colla, E. Linking the Endoplasmic Reticulum to Parkinson’s Disease and Alpha-Synucleinopathy. Front Neurosci 2019, 13, doi:10.3389/fnins.2019.00560.

38. Smith, W.W.; Jiang, H.; Pei, Z.; Tanaka, Y.; Morita, H.; Sawa, A.; Dawson, V.L.; Dawson, T.M.; Ross, C.A. Endoplasmic Reticulum Stress and Mitochondrial Cell Death Pathways Mediate A53T Mutant Alpha-Synuclein-Induced Toxicity. Hum Mol Genet 2005, 14, doi:10.1093/hmg/ddi396.

39. Heman-Ackah, S.M.; Manzano, R.; Hoozemans, J.J.M.; Scheper, W.; Flynn, R.; Haerty, W.; Cowley, S.A.; Bassett, A.R.; Wood, M.J.A. Alpha-Synuclein Induces the Unfolded Protein Response in Parkinson’s Disease SNCA Triplication IPSC-Derived Neurons. Hum Mol Genet 2017, 26, doi:10.1093/hmg/ddx331.

40. Smith, J.K. Osteoclasts and Microgravity. Life 2020, Vol. 10, Page 207 2020, 10, 207, doi:10.3390/LIFE10090207.

41. Buoite Stella, A.; Ajčević, M.; Furlanis, G.; Manganotti, P. Neurophysiological Adaptations to Spaceflight and Simulated Microgravity. Clin Neurophysiol 2021, 132, 498–504, doi:10.1016/J.CLINPH.2020.11.033.

42. Juhl, O.J.; Buettmann, E.G.; Friedman, M.A.; DeNapoli, R.C.; Hoppock, G.A.; Donahue, H.J. Update on the Effects of Microgravity on the Musculoskeletal System. npj Microgravity 2021 7:1 2021, 7, 1–15, doi:10.1038/s41526-021-00158-4.

43. Wu, X.T.; Yang, X.; Tian, R.; Li, Y.H.; Wang, C.Y.; Fan, Y.B.; Sun, L.W. Cells Respond to Space Microgravity through Cytoskeleton Reorganization. FASEB Journal 2022, 36, doi:10.1096/FJ.202101140R.

44. Nguyen, H.P.; Tran, P.H.; Kim, K.S.; Yang, S.G. The Effects of Real and Simulated Microgravity on Cellular Mitochondrial Function. npj Microgravity 2021 7:1 2021, 7, 1–11, doi:10.1038/s41526-021-00171-7.

45. Amselem, S. Remote Controlled Autonomous Microgravity Lab Platforms for Drug Research in Space. Pharm Res 2019, 36, 1–15, doi:10.1007/S11095-019-2703-7/FIGURES/8.

46. Roy-O’Reilly, M.; Mulavara, A.; Williams, T. A Review of Alterations to the Brain during Spaceflight and the Potential Relevance to Crew in Long-Duration Space Exploration. NPJ Microgravity 2021, 7, doi:10.1038/S41526-021-00133-Z.

47. Stavnichuk, M.; Mikolajewicz, N.; Corlett, T.; Morris, M.; Komarova, S. V. A Systematic Review and Meta-Analysis of Bone Loss in Space Travelers. NPJ Microgravity 2020, 6, doi:10.1038/S41526-020-0103-2.

48. Frippiat, J.P.; Crucian, B.E.; de Quervain, D.J.F.; Grimm, D.; Montano, N.; Praun, S.; Roozendaal, B.; Schelling, G.; Thiel, M.; Ullrich, O.;, et al. Towards Human Exploration of Space: The THESEUS Review Series on Immunology Research Priorities. NPJ Microgravity 2016, 2, doi:10.1038/NPJMGRAV.2016.40.

49. Tauber, S.; Hauschild, S.; Paulsen, K.; Gutewort, A.; Raig, C.; Hürlimann, E.; Biskup, J.; Philpot, C.; Lier, H.; Engelmann, F.;, et al. Signal Transduction in Primary Human T Lymphocytes in Altered Gravity During Parabolic Flight and Clinostat Experiments. Cellular Physiology and Biochemistry 2015, 35, 1034–1051, doi:10.1159/000373930.

50. Zhang, Y.; Wang, H.; Lai, C.; Wang, L.; Deng, Y. Comparative Proteomic Analysis of Human SH-SY5Y Neuroblastoma Cells under Simulated Microgravity. Astrobiology 2013, 13, 143–150, doi:10.1089/AST.2012.0822/ASSET/IMAGES/LARGE/FIGURE7.JPEG.

51. Rampoldi, A.; Forghani, P.; Li, D.; Hwang, H.; Armand, L.C.; Fite, J.; Boland, G.; Maxwell, J.; Maher, K.; Xu, C. Space Microgravity Improves Proliferation of Human IPSC-Derived Cardiomyocytes. Stem Cell Reports 2022, 17, doi:10.1016/j.stemcr.2022.08.007.

52. Striebel, J.; Kalinski, L.; Sturm, M.; Drouvé, N.; Peters, S.; Lichterfeld, Y.; Habibey, R.; Hauslage, J.; El Sheikh, S.; Busskamp, V.;, et al. Human Neural Network Activity Reacts to Gravity Changes in Vitro. Front Neurosci 2023, 17, doi:10.3389/fnins.2023.1085282.

53. Wang, J.; Zhang, J.; Bai, S.; Wang, G.; Mu, L.; Sun, B.; Wang, D.; Kong, Q.; Liu, Y.; Yao, X.;, et al. Simulated Microgravity Promotes Cellular Senescence via Oxidant Stress in Rat PC12 Cells. Neurochem Int 2009, 55, 710–716, doi:10.1016/J.NEUINT.2009.07.002.

54. Pala, R.; Cruciani, S.; Manca, A.; Garroni, G.; EL Faqir, M.A.; Lentini, V.; Capobianco, G.; Pantaleo, A.; Maioli, M. Mesenchymal Stem Cell Behavior under Microgravity: From Stress Response to a Premature Senescence. Int J Mol Sci 2023, 24, doi:10.3390/IJMS24097753.

55. Di Giulio, C. Do We Age Faster in Absence of Gravity? Front Physiol 2013, 4, doi:10.3389/FPHYS.2013.00134.

56. Fukuda, A.; … V. de L.C.-L.S. in S.; 2021, undefined Simulated Microgravity Accelerates Aging in Saccharomyces Cerevisiae. Elsevier.

57. Dinarelli, S.; Longo, G.; Dietler, G.; Francioso, A.; Mosca, L.; Pannitteri, G.; Boumis, G.; Bellelli, A.; Girasole, & M. Erythrocyte’s Aging in Microgravity Highlights How Environmental Stimuli Shape Metabolism and Morphology. Springer 2018, 8, 5277, doi:10.1038/s41598-018-22870-0.

58. Ratushnyy, A.Y.; Buravkova, L.B. Microgravity Effects and Aging Physiology: Similar Changes or Common Mechanisms? Biochemistry (Moscow) 2023, 88, 1763–1777.

59. Prasad, B.; Grimm, D.; Strauch, S.M.; Erzinger, G.S.; Corydon, T.J.; Lebert, M.; Magnusson, N.E.; Infanger, M.; Richter, P.; Krüger, M. Influence of Microgravity on Apoptosis in Cells, Tissues, and Other Systems in Vivo and in Vitro. Int J Mol Sci 2020, 21, doi:10.3390/ijms21249373.

60. Uras, G.; Li, X.; Manca, A.; Pantaleo, A.; Bo, M.; Xu, J.; Allen, S.; Zhu, Z. Development of P-Tau Differentiated Cell Model of Alzheimer’s Disease to Screen Novel Acetylcholinesterase Inhibitors. Int J Mol Sci 2022, 23, 14794, doi:10.3390/IJMS232314794/S1.

61. Xicoy, H.; Wieringa, B.; Martens, G.J.M. The SH-SY5Y Cell Line in Parkinson’s Disease Research: A Systematic Review. Mol Neurodegener 2017, 12.

62. Lucas-Del-Pozo, #; Sara; Uras,; #; Giuseppe; Fierli, ; Federico; Lentini, ; Veronica; Koletsi, ; Sofia; et al. Reduction of A-Synuclein Aggregates by PIKfyve Inhibition via TFEB-Mediated Lysosomal Biogenesis in a Parkinson Disease Model. bioRxiv 2023, 2023.10.23.563557, doi:10.1101/2023.10.23.563557.

63. Terry-Kantor, E.; Tripathi, A.; Imberdis, T.; Lavoie, Z.M.; Ho, G.P.H.; Selkoe, D.; Fanning, S.; Ramalingam, N.; Dettmer, U. Rapid Alpha-Synuclein Toxicity in a Neural Cell Model and Its Rescue by a Stearoyl-CoA Desaturase Inhibitor. International Journal of Molecular Sciences 2020, Vol. 21, Page 5193 2020, 21, 5193, doi:10.3390/IJMS21155193.

64. Wuest, S.L.; Richard, S.; Kopp, S.; Grimm, D.; Egli, M. Simulated Microgravity: Critical Review on the Use of Random Positioning Machines for Mammalian Cell Culture. Biomed Res Int 2015, 2015.

65. Bradshaw, A. V.; Campbell, P.; Schapira, A.H.V.; Morris, H.R.; Taanman, J.W. The PINK1-Parkin Mitophagy Signalling Pathway Is Not Functional in Peripheral Blood Mononuclear Cells. PLoS One 2021, 16, doi:10.1371/journal.pone.0259903.

66. Uras, G. Evaluation of AChE Inhibitors and Dual AChE/GSK3-β Inhibitor as Alzheimer’s Disease Treatment;

67. Costa, C.A. da; Manaa, W. El; Duplan, E.; Checler, F. The Endoplasmic Reticulum Stress/Unfolded Protein Response and Their Contributions to Parkinson’s Disease Physiopathology. Cells 2020, Vol. 9, Page 2495 2020, 9, 2495, doi:10.3390/CELLS9112495.

68. Colla, E. Linking the Endoplasmic Reticulum to Parkinson’s Disease and Alpha-Synucleinopathy. Front Neurosci 2019, 13, 461767, doi:10.3389/FNINS.2019.00560/BIBTEX.

69. Kawahata, I.; Finkelstein, D.I.; Fukunaga, K. Pathogenic Impact of α-Synuclein Phosphorylation and Its Kinases in α-Synucleinopathies. International Journal of Molecular Sciences 2022, Vol. 23, Page 6216 2022, 23, 6216, doi:10.3390/IJMS23116216.

70. Stewart, T.; Sossi, V.; Aasly, J.O.; Wszolek, Z.K.; Uitti, R.J.; Hasegawa, K.; Yokoyama, T.; Zabetian, C.P.; Leverenz, J.B.; Stoessl, A.J.;, et al. Phosphorylated α-Synuclein in Parkinson’s Disease: Correlation Depends on Disease Severity. Acta Neuropathol Commun 2015, 3, 7, doi:10.1186/S40478-015-0185-3/FIGURES/3.

71. Fujiwara, H.; Hasegawa, M.; Dohmae, N.; Kawashima, A.; Masliah, E.; Goldberg, M.S.; Shen, J.; Takio, K.; Iwatsubo, T. α-Synuclein Is Phosphorylated in Synucleinopathy Lesions. Nat Cell Biol 2002, 4, doi:10.1038/ncb748.

72. Mbefo, M.K.; Fares, M.B.; Paleologou, K.; Oueslati, A.; Yin, G.; Tenreiro, S.; Pinto, M.; Outeiro, T.; Zweckstetter, M.; Masliah, E.;, et al. Parkinson Disease Mutant E46K Enhances α-Synuclein Phosphorylation in Mammalian Cell Lines, in Yeast, and in Vivo. J Biol Chem 2015, 290, 9412, doi:10.1074/JBC.M114.610774.

73. Esterbauer, H.; Schaur, R.J.; Zollner, H. Chemistry and Biochemistry of 4-Hydroxynonenal, Malonaldehyde and Related Aldehydes. Free Radic Biol Med 1991, 11.

74. Marnett, L.J. Oxy Radicals, Lipid Peroxidation and DNA Damage. Toxicology 2002, 181–182, doi:10.1016/S0300-483X(02)00448-1.

75. Zhang, Y.; Wang, H.; Lai, C.; Wang, L.; Deng, Y. Comparative Proteomic Analysis of Human SH-SY5Y Neuroblastoma Cells under Simulated Microgravity. https://home.liebertpub.com/ast 2013, 13, 143–150, doi:10.1089/AST.2012.0822.

76. Colla, E.; Coune, P.; Liu, Y.; Pletnikova, O.; Troncoso, J.C.; Iwatsubo, T.; Schneider, B.L.; Lee, M.K. Endoplasmic Reticulum Stress Is Important for the Manifestations of α-Synucleinopathy in Vivo. Journal of Neuroscience 2012, 32, doi:10.1523/JNEUROSCI.5367-11.2012.

77. Kim, S.; Kim, D.K.; Jeong, S.; Lee, J. The Common Cellular Events in the Neurodegenerative Diseases and the Associated Role of Endoplasmic Reticulum Stress. Int J Mol Sci 2022, 23.

78. Karvandi, M.S.; Sheikhzadeh Hesari, F.; Aref, A.R.; Mahdavi, M. The Neuroprotective Effects of Targeting Key Factors of Neuronal Cell Death in Neurodegenerative Diseases: The Role of ER Stress, Oxidative Stress, and Neuroinflammation. Front Cell Neurosci 2023, 17.

79. Chou, S.M.; Yen, Y.H.; Yuan, F.; Zhang, S.C.; Chong, C.M. Neuronal Senescence in the Aged Brain. Aging Dis 2023, 14.

80. Byers, B.; Cord, B.; Nguyen, H.N.; Schüle, B.; Fenno, L.; Lee, P.C.; Deisseroth, K.; Langston, J.W.; Pera, R.R.; Palmer, T.D. SNCA Triplication Parkinson’s Patient’s IPSC-Derived DA Neurons Accumulate α-Synuclein and Are Susceptible to Oxidative Stress. PLoS One 2011, 6, e26159, doi:10.1371/JOURNAL.PONE.0026159.

81. Nguyen, H.N.; Byers, B.; Cord, B.; Shcheglovitov, A.; Byrne, J.; Gujar, P.; Kee, K.; Schüle, B.; Dolmetsch, R.E.; Langston, W.;, et al. LRRK2 Mutant IPSC-Derived DA Neurons Demonstrate Increased Susceptibility to Oxidative Stress. Cell Stem Cell 2011, 8, 267–280, doi:10.1016/J.STEM.2011.01.013.

82. Manis, C.; Manca, A.; Murgia, A.; Uras, G.; Caboni, P.; Congiu, T.; Faa, G.; Pantaleo, A.; Cao, G. Understanding the Behaviour of Human Cell Types under Simulated Microgravity Conditions: The Case of Erythrocytes. Int J Mol Sci 2022, 23, doi:10.3390/ijms23126876.

83. Yakunin, E.; Kisos, H.; Kulik, W.; Grigoletto, J.; Wanders, R.J.A.; Sharon, R. The Regulation of Catalase Activity by PPAR γ Is Affected by α-Synuclein. Ann Clin Transl Neurol 2014, 1, 145–159, doi:10.1002/ACN3.38.

